# Controlling insulin resistance through modulation of interplay between mediators of cellular senescence: A mathematical study

**DOI:** 10.1101/269738

**Authors:** Chakit Arora

**Author notes:** presently at Center for Computational Biology, Indraprastha Institute of Information Technology, Okhla Phase III, Delhi-110020, India.

## Abstract

Obesity, metabolic syndrome and premature ageing form a hugely researched and discussed area of interest these days. In the pathology of this cluster of conditions, adipose tissue is gaining attention as a major playground for interplay between metabolic stress, inflammation and accelerated ageing, and not merely being an energy storage tank. Drastic elevation in the levels of reactive oxygen species, due to lipid overload and excessive lipolysis, causes genotoxic damage such as shortening of telomeres (an indicator of accelerated cell ageing), increased mRNA and protein expression for p53, p21, TNF–*α*, IL-6 (interleukin 6), impaired insulin mediated glucose uptake, and decreased TERT mRNA expression. The increase of p53 in adipocytes is deleterious, since elevated p53 results in pre-mature ageing of fat tissues which secrete pro-inflammatory cytokines thereby contributing to insulin resistance. But inhibition of p53 as a therapeutic target, as suggested by many previous studies, could result in developing high risks of cancer. The association between hyperlipidemic/hyperglycaemic stress, premature growth arrest and insulin resistance, thus, forms an interesting premise for searching targets and designing interventions for therapy of metabolic syndrome, type 2 diabetes mellitus etc. We developed a mathematical study involving a 5D ODE model, which revealed crucial parametric conditions governing p53 dynamics in the case of metabolic stress-induced cellular senescence, and shed light on potential strategies to reduce pro-inflammatory cytokine levels which exacerbate insulin resistance through premature cellular ageing.

**Highlights:**

1. p53 oscillations signify DNA repair, and persistent stress causes prolonged surge in p53 levels inducing cellular senescence.
2. p53-induced cellular senescence promotes inflammation and enhances progression of insulin resistance.
3. Regulation of Mdmx and Akt can be a strategy to rejuvenate ageing cells through management of IL-6 dynamics.

## 1 Introduction

Diabetes mellitus, obesity and associated metabolic disorders are reaching pandemic proportions in their incidence [1, 2, 3]. These are marked by metabolic disturbances such as hyperglycemia, dyslipidemia etc. that result in molecular level derangements, such as generation of reactive oxygen species and inflammatory markers. Oxidative stress targets and damages an array of biomolecules, DNA being one of them, where damage leads to either activation of repair mechanism, or growth arrest/apoptosis [7, 8]. This has been observed in adipose tissue, where increased levels of ROS and excessive lipolysis are followed by premature cellular senescence and elevated pro-inflammatory marker profile [9]. This pro-inflammatory state eventually leads to development of insulin resistance, and is a hallmark of metabolic syndrome.

One of the principal factors that confer senescence-like phenotype to the tissue under conditions of metabolic dysregulation is the classical tumour-suppressor protein p53 [9]. DNA damage due to oxidative stress leads to activation and elevated levels of p53, a master regulator of cell cycle, which further activates p21 and eventually leads to higher levels of the pro-inflammatory cytokine IL-6 [8]. Elevation in IL-6 levels has been shown to be associated with development of insulin resistance, the principal feature of type 2 diabetes mellitus [38]. This pathway has been validated experimentally where insulin resistance in adipose tissue was found to be associated with high expression of p53 [10].

p53 is one of the most popularly studied genes owing to its crucial role as a key player in prevention of development of cancer [60, 61, 62, 63]. Various mathematical models have been proposed for studying its behavioral dynamics, to explain the experimental results, as well as to gain a insight into its relationship with the other associated elements for the desired action [39, 4, 5, 6, 12]. One such experimentally observed phenomenon is the oscillatory levels of p53 protein as reported in [12], which corresponds to DNA repair induced upon encounter of the cell with a genotoxic stress.

In order to have a molecular network resulting in oscillatory behaviour, we are restricted by some standard mathematically established topologies and constraints in kinetics [40, 41, 42]. The molecular network then could be a negative feedback loop with a delay or a combination of positive feedback and negative feedback loops [39]. A commonly used trick is to take the product of the sign of edges, a negative sign as a result would imply that the network can exhibit oscillations (necessary but not sufficient).

In the present paper we took a simplistic network which forms a long negative feedback loop as given in [12], comprising three key players p53, Mdm2 and Mdmx. However they took an intermediate subsidiary I to explain their model and results, which has been replaced here by the homologous protein called Mdmx which inhibits p53 activity and stabilises Mdm2 [13]. We used their 5-D model concept to build our three-dimensional model. The 3-D model thus obtained by modification was then extended to a 5-D non autonomous ode model to explain the roles of p21, and IL6 or Akt. Our study showed that there exist parametric conditions such that one may have normal IL6 levels without altering p53 dynamics under exposure to stress.

## 2 The Model

The concept of model is schematically described in figure 1. Here the input represents (oxidative) stress, output 1 shows the dynamics of p53 levels and output 2 represents IL6 dynamics. So, essentially we have divided our model into 2 parts namely part I and part II. These are explained sequentially in following manner.

**Figure 1:**
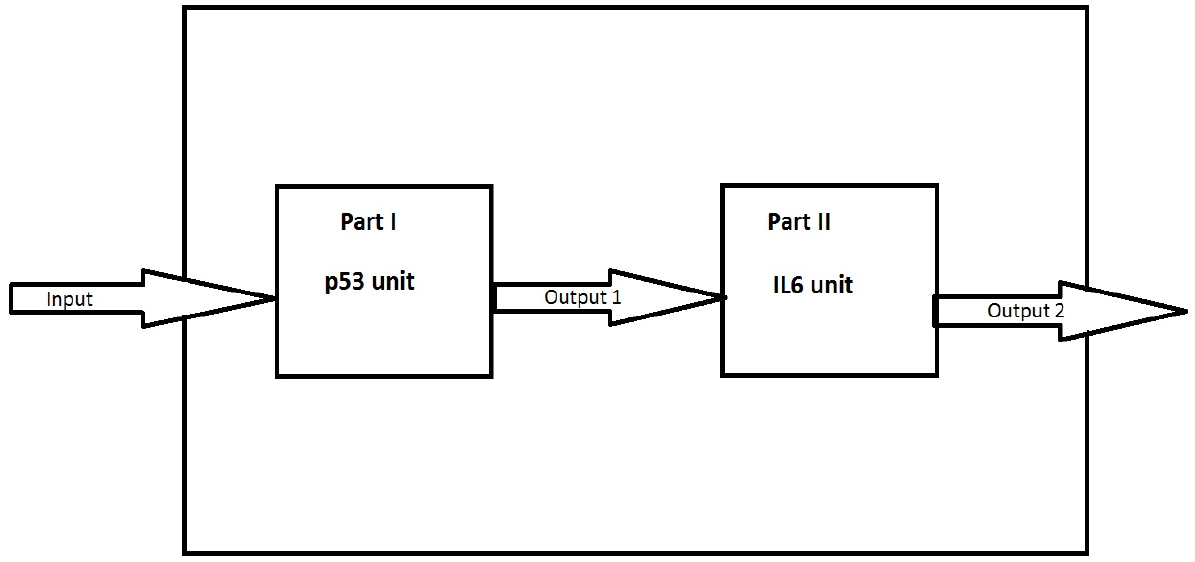
The figure is schematic block diagram of the model. The complete model is represented by the combination of two sub units : part I representing a 3-D network comprising p53,Mdmx and Mdm2; part II is a 2D network following unit I comprising of IL6 and p21. The input here is the stress signal acting on unit I. The two outputs-output 1 and output 2-represent the output of unit I i.e p53 and the output 2 reads out the levels of IL6 and hence Akt.

### 2.1 Part I : The p53 unit

This part of the model contains a 3-D non autonomous system of ode’s obtained from a 5-D model proposed in [12]. *y*_1_(*t*), *y*_2_(*t*), *y*_3_(*t*) are the concentrations of p53 protein, Mdm2 protein and Mdmx protein respectively. It was noted in previous studies that p53 levels were elevated during stressed conditions, the opposite of which is seen during non stressed conditions. Also Mdm2 has been widely recognised as crucial regulator of p53 as its absence leads to excess activity of p53 causing embryonic death [15] while excessive Mdm2 expression leads to inhibition of p53 and thereby promotes cancer [15, 16, 17, 18]. Mdm2 was found to negatively regulate p53 by acting as an ubiquitin ligase or by proteasomal degradation [19, 20, 21, 22]. This fact is portrayed in Eqn. 1 by the term ‘−*d*_1_*y*_1_*y*_2_’ where *d*_1_ represents rate of Mdm2 dependent degradation of p53. The first term *s*_1_ denotes synthesis rate of p53 protein. The last term of Eqn. 1 represents effect of input on p53 degradation, more specifically it shows that during high stress (t=0) the Mdm2 dependent degradation of p53 is lowest and as stress fades away it reaches its maximum value viz. ‘−*d*_1_*y*_1_*y*_2_’. We have taken the term ‘*e*^−*σt*^’ to represent the oxidative stress conditions (input1) as a result of accumulation of ROS due to lipid breakdown in the adipose tissue. This factor represents a signal which is maximum at t=0 implying the time at which the network faces a stress pulse. The stress pulse eventually fades away as defined by *σ*.

Eqn. (2) shows the rate equation for Mdm2 protein. *p*_1_ represents rate of synthesis of Mdm2 protein independent of p53 [23, 24]. The last term represents Mdm2 degradation [25]. The second term shows stabilisation of Mdm2 by Mdmx [26, 27]. Eqn. (3) represents rate of Mdmx protein, where the second term shows the degradation of Mdmx independent of Mdm2 [28] and the first term shows the enhancement of Mdmx protein production by p53 [29] and input stress while Mdm2 represses it by binding to p53 and reducing its transcriptional activity [43]. The sub-model I is written as:

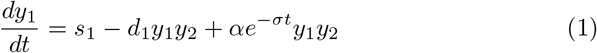

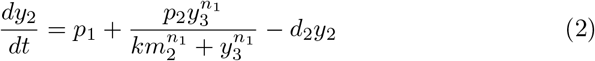

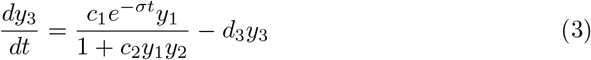

For our mathematical study we assumed that the system is in a steady state initially before the action of stress. During stressed condition, there is a change in the state of the variables. But, when the stress fades away to zero, the system again goes back to the same initial state. Steady state here is the state (*y*_1_*, *y*_2_*, *y*_3_*) at *t* → ∞ i.e when the exponential term is zero. This asymptotic limit of model is equivalent to a scenario where the system didn’t face the stress. In essence we have a system at a stable steady state which faces an input, reacts to it dynamically and then comes back to the stable state after the input is removed (see section 3.1.1 for detailed analysis). This hypothesis is validated via reproduction of results as in [12] (see figure 3).

### 2.2 Part II : The IL6 unit

There is a significant amount of literature which links p21 downstream to p53 which plays a regluatory role during progression to senescence when transcribed by p53 in response to DNA damage [30]. The p21 protein then further activates a series of downstream mediators ultimately forcing the cell into cellular senescence as a protective measure against stress. Adipocytes subjected to oxidative stress also showed increased expression of pro-inflammatory cytokines TNF-α and IL-6 [8]. It has been reported that increased secretion of pro-inflammatory cytokines by adipose tissue exacerbates insulin resistance in people with metabolic disorders [31, 32]. We have singled out IL-6 from amongst all the cytokines associated with the senescent phenotype due to its strong association with insulin resistance with as well as independent of obesity, and more importantly its endocrine nature unlike other cytokines such as TNF-a that are secreted by a limited number of types of cells and act in a paracrine fashion [53] . In this unit of the model we tried to include this mechanism of p21 transcription by p53 as shown in Eqn. (4) and the synthesis of IL6 via senescence pathways as shown in Eqn. (5). It is also been found that the expression of IL6 causes an increase in expression of Akt which negatively controls p21 through phosphorylation [33, 34].

Since there is a linear relationship between IL6 and Akt [47], so, an extra equation for Akt is not required and the concentration of Akt is taken to be directly proportional to IL6. The sub-model II is written as:

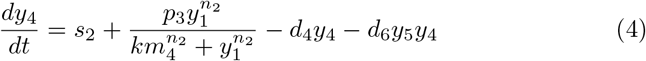

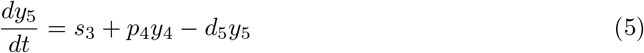

The variables *y*_4_(*t*) and *y*_5_(*t*) represent the concentration of protein p21 and cytokine IL6 respectively. Eqn. (4) gives the rate of p21 production where *s*_2_ is its synthesis rate independent of p53 [35, 36]. *p*_3_ in the second hills function type term of Eqn. (4) is the p53 dependent rate of p21 transcription. This *p*_3_ may define the condition for which p21 transcription takes place. *d*_4_ represent degradation of p21 protein where *d*_6_ is the rate dependent on degradation of p21 due to Akt. Eqn. (5) gives the rate of IL6 synthesis, which then proportionally activates Akt (not explicitly mentioned in the model). *s*_3_ is the basal production rate independent of p53 and p21 mechanism [37] while *p*_4_ is the rate of IL6 synthesis via senescent pathway activated by p21. *d*_5_ is the degradation rate of IL6 through other mechanisms [37].

### 2.3 The Complete Model

Our complete model (6) consists of the combination of both the units above connected in the fashion as depicted by figure 1.

This model explains on a dynamical basis the tight link that exists between p53 function as a tumor supressor and insulin resistance when adipocytes are under high oxidative stress. The following 5-D non autonomous ode system is represented in detail in figure 2.

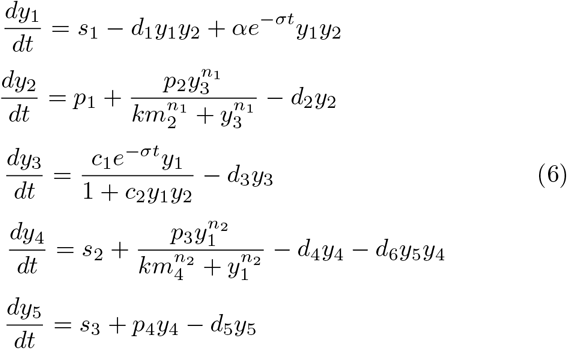

**Figure 2:**
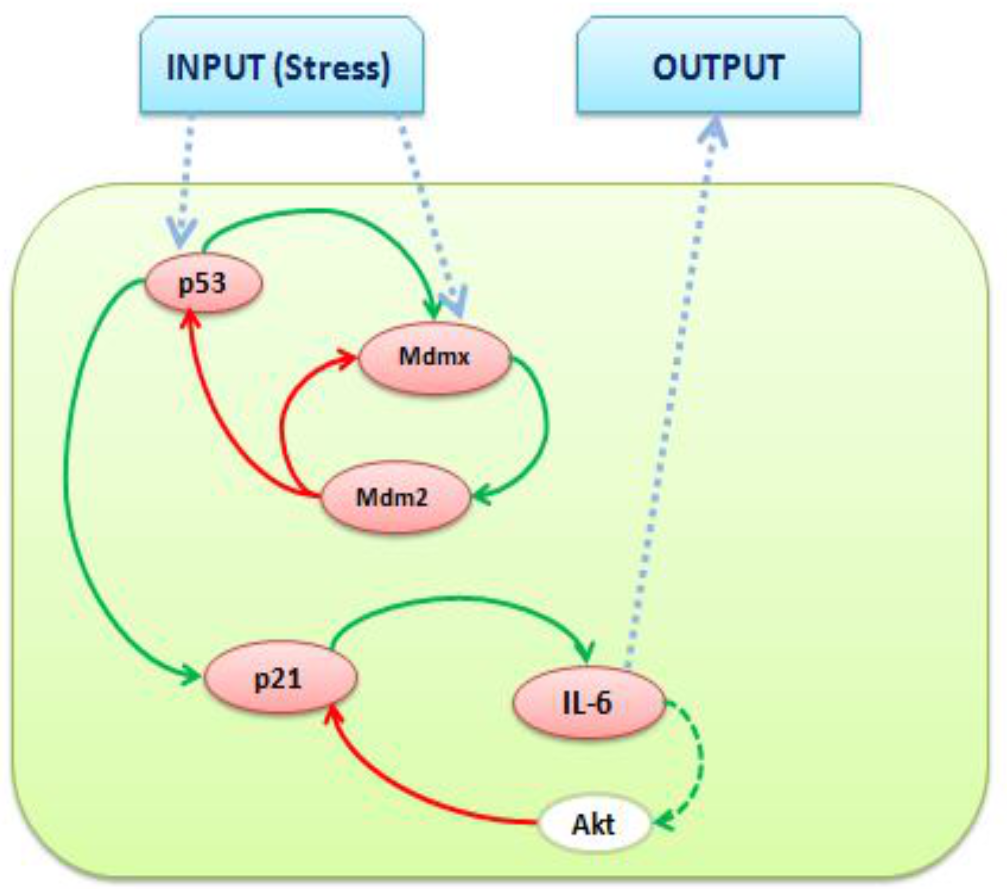
The complete model : The diagram above shows the network structure of our model. The green arrows mean positive effect of an element on the tail of arrow to the element on the head of arrow. The red arrows represents a negative effect in the same manner.

## 3 Results

### 3.1 Mathematical Analysis

#### 3.1.1 The p53 Unit

The main hypothesis used here is that the system comes back to its initial state after the stress fades away. Suppose we have a non autonomous system with an input I(t), which vanishes after a certain amount of time, and whenever the system encounters this stimulus I(t), it reacts accordingly. But it is to be noted that the absence of any stimulus in first place and fading away of the stimulus after a certain amount of time, are mathematically equivalent conditions (i.e *I*(*t*) = 0). Thus it is essential for the system to revert back to its initial state, after the stimulus vanishes, to justify this equivalence. One can use this hypothesis in the following manner:

a one dimensional non autonomous ode system can be written as 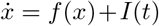 where *f*(*x*) is the time independent part and *I*(*t*) is the input stimulus to the system. For a case when I is not present, one can then find fixed points for the equation 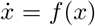. Assuming that *x*\* represents the stable fixed point for this equation, where *x*\* has a constant value for a fixed set of the parameter values. This *x*\* can be made equal to the initial value *x*_0_ of the system with some choice of parameters. Thus we can define a system with initial value *x*(0) = *x*_0_ and a stable fixed point *x*\* = *x*_0_ which means that the system dynamics after initiating from *x*_0_ following some trajectories will come back to *x*_0_, and will remain at *x*_0_ for all the future time.

Now keeping the above design intact, we introduce a stimulus I(t) to this system where *I*(*t*) > 0 for *t* ∈ (*t*_1_, *t*_2_) and zero otherwise, and observe the ‘reaction’ of the system. So we have actually designed a system which starts with some initial ‘unperturbed’ state, goes to a ‘perturbed’ state when fed with a stimulus, and comes back to the ‘initial’ state after the stimulus is removed.

Following the above hypothesis in our model, we take the asymptotic limit *t* → ∞ where our sub-model I reduces to the following set of ode’s:

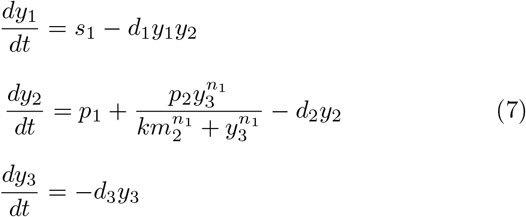

One could then perform a fixed point analysis on system (7) to show that at such an asymptotic limit the system reaches the stable fixed point

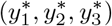

where

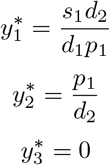

Now, in order to obtain trajectory which has the same initial state and final state, we set the following conditions:

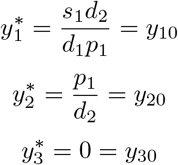

where (*y*_10_, *y*_20_, *y*_30_) is the initial state of the system. So, the steady state value could be made equal to the initial state by using the parameters *p*_1_, *d*_2_, *s*_1_ and *d*_1_. Following this, we studied the parameter-dependent model sensitivity in order to resolve the stress input as well as mimic the previously observed phenomena. To validate our reduced 3-D model with the model given in [12], we reproduced their results using table 1, where most of the parameter values were taken from their study (see figure 3).

**Table 1:** Basal values for the sub-model I parameters for oscillation in p53 levels (DNA repair) [12]. Here initial conditions are *y*_10_ = 5.3191, *y*_20_ = 0.047 and *y*_30_ = 2.

| Parameter | Description | Value |
| --- | --- | --- |
| $s_1$ | Basal production rate of p53 | 0.5 |
| $d_1$ | Degradation of p53 via Mdm2 | 2 |
| $p_1$ | Basal production rate of Mdm2 | 0.00235 |
| $p_2$ | Mdmx dependent production rate of Mdm2 | 0.03 |
| $d_2$ | Degradation of Mdm2 | 0.05 |
| $c_1$ | Input dependent production rate of Mdmx | 0.09 |
| $c_2$ | Mdm2 dependent degradation of Mdmx | 0.01 |
| $d_3$ | Degradation rate of Mdmx | 0.09 |
| $\alpha$ | Rate of stress dependent p53 enhancement | 1.93 |
| $\sigma$ | Resolution rate of stress signal | 0.0001 |
| $km_2$ | Michealis constt in production of Mdm2 via Mdmx | 25 |
| $n_1$ | Hill's coefficient in production of Mdm2 via Mdmx | 50 |

**Figure 3:**
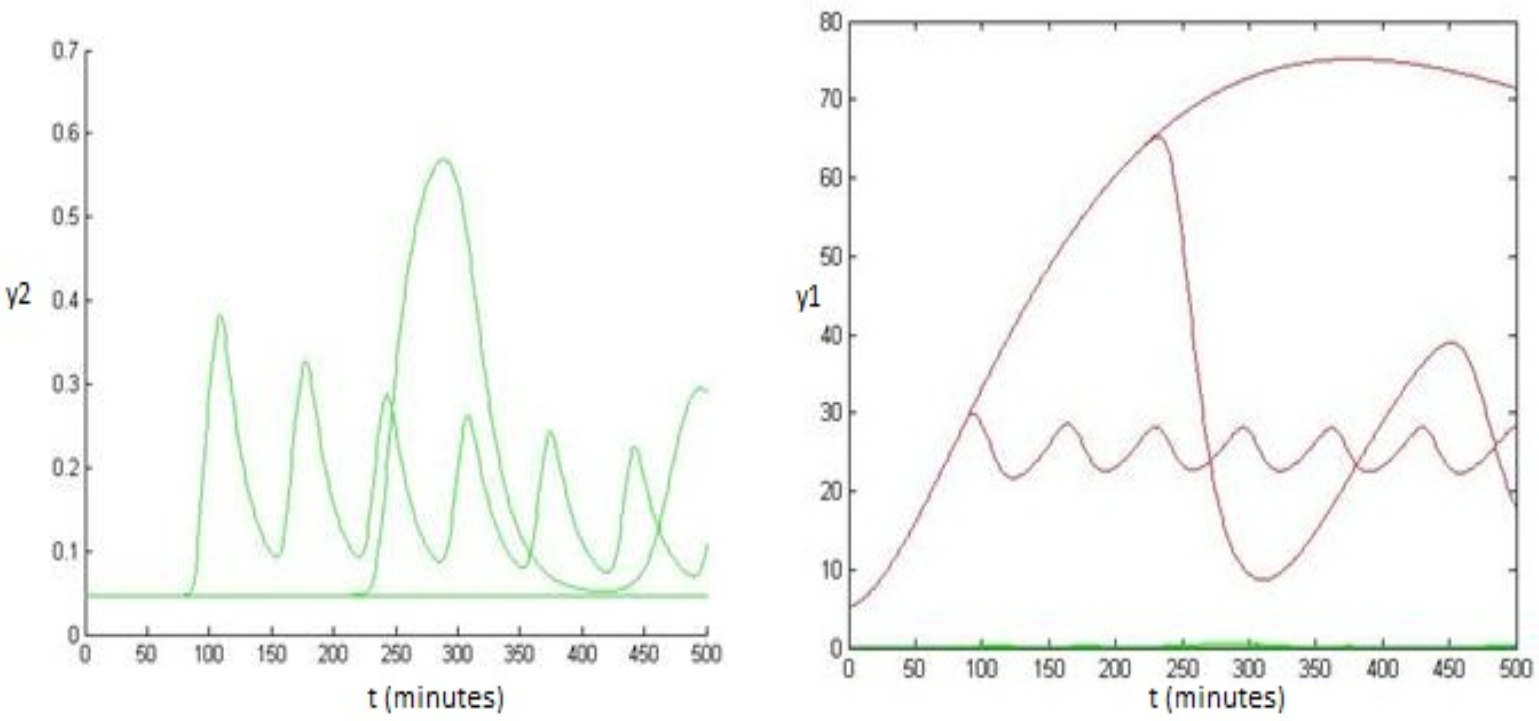
The figure is a reproduction of alon’s results [12] where we used our modified model and kept the model parameters as earlier in addition to which we set *α* = 1.93 and *σ* = 0.0001. Red represents p53 and green represents Mdm2.

We then took table 1 as our basal set of parameters, where oscillations (DNA repair) take place, and performed a sensitivity analysis as shown in figure 4 to find out the sensitive parameters pertaining to oscillatory state of the system.

**Figure 4:**
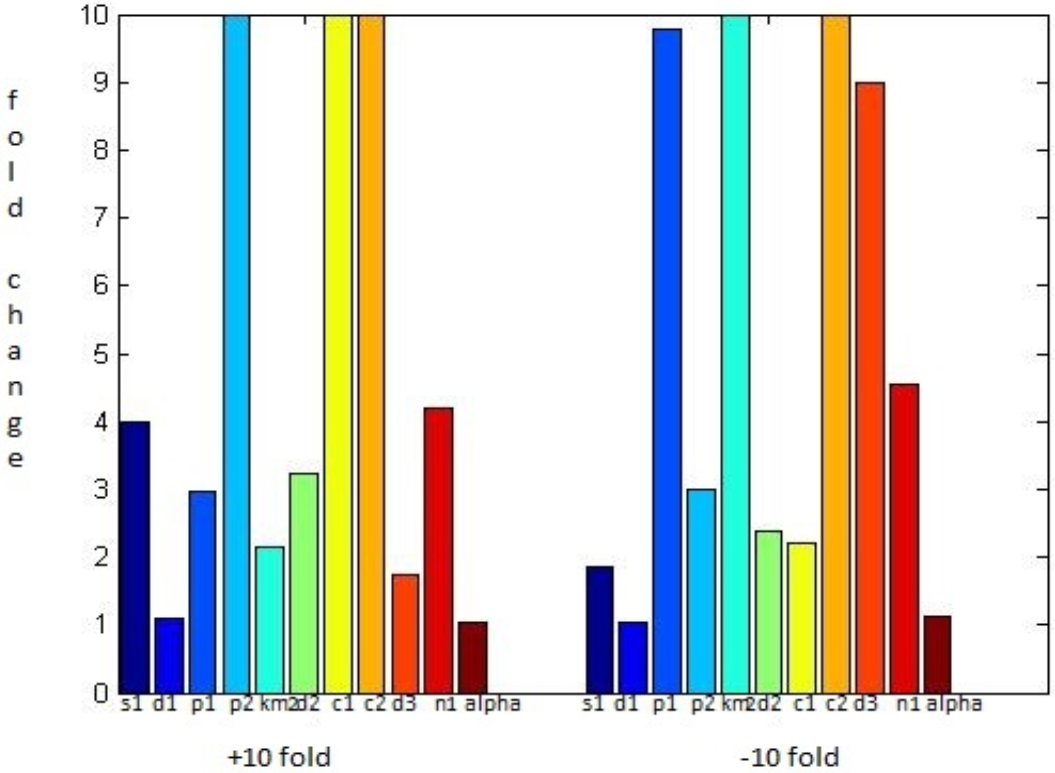
The diagram represents the sensitivity analysis of the 3-D model in the p53 unit. A bar here represent the fold change values of a parameter around the basal value such that oscillations take place, while keeping other parameters fixed. It is evident that *d*_1_, *s*_1_ and *α* are the most sensitive parameters amongst all.

#### 3.1.2 The IL6 Unit

Let us take a situation when the transcription rate of p21 due to p53 is low i.e

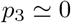

which is equivalent to the situation where the senescence pathway is not activated. Thus our sub-model II gets reduced to the following set of odes:

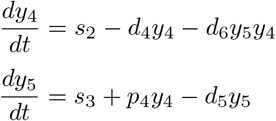

The fixed point (*y*_4_*, *y*_5_*) for the above system is found, where:

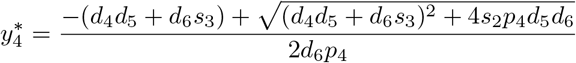

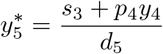

for a biologically feasible solution to exist, we have

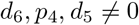

Now if we take the parameters

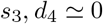

representing the fact that production of IL6 through other independent processes is negligible and degradation of p21 through an Akt independent way is also insignificant. We then have the following results:

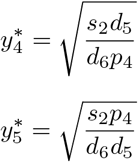

Using these inferences we fixed the parameter values in order to mimic the experimental values of IL6 when there is no stress or senescence like conditions.

#### 3.1.3 The Complete Model : Positivity and Boundedness

The complete model (6) can be written in the form:

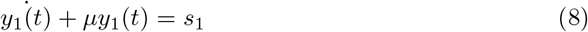

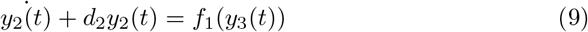

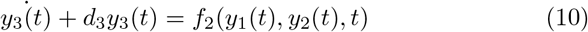

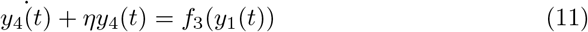

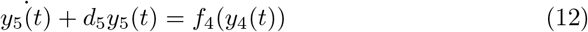

where

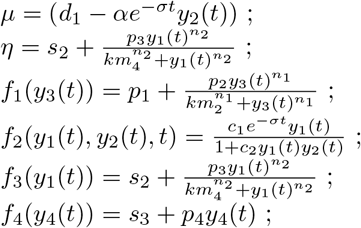

for simplification purposes we will use shorthand notations *f*_1_, *f*_2_, *f*_3_ and *f*_4_ to denote the above functions. Using this notation and keeping in mind that all parameters are positive, we analysed the system for positivity and boundedness conditions.

##### Positivity

We perform the following stepwise analysis for positivity conditions of the system:

1. From Eqn. (10) above and also noticing the fact that *f*_2_ vanishes at *y*_1_(*t*) = 0 or as *t* → ∞. We obtain the solution for *y*_3_(*t*) as

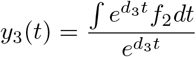

thus *y*_3_(*t*) ≥ 0 since the numerator is positive or zero depending on *f*_2_.
2. From Eqn. (9) and using condition 1 above we have *f*_1_ > 0 for every *y*_3_(*t*). The solution for *y*_2_(*t*) is found to be

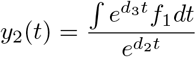

which is positive definite since the numerator is always positive.
3. From Eqn. (8) and using the above conditions we have

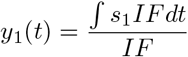

where

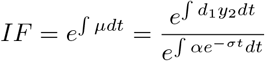

and hence *y*_1_(*t*) > 0 since *IF* > 0 for positive values of t.
4. From Eqn. (11) and using the fact that *η*, *f*_3_ > 0 the solution

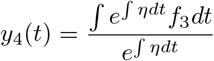

is always positive.
5. From Eqn. (12) and using result above the solution

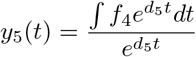

is always positive.

##### Boundedness

We will use the terminology and results of the above section to show that the solutions are bounded for any time ‘t’ within a range whose lower bounds were found as *y*_1_(*t*), *y*_2_(*t*), *y*_4_(*t*), *y*_5_(*t*) > 0 and *y*_3_(*t*) ≥ 0. This analysis is shown in the following steps:

1. From Eqn. (9) one can write : 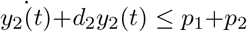 since *max*(*f*_1_) = *p*_1_+*p*_2_ therefore the solution is bounded by the condition

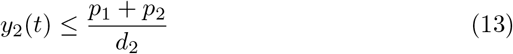
2. From Eqn. (8) we can have a boundedness condition for *y*_1_(*t*) in a similar fashion as:
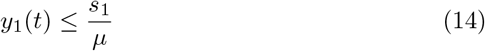

where

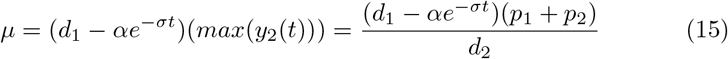

since 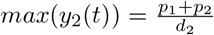 from (13). It should be noted that this boundedness condition holds true only when

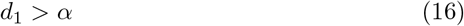
3. From Eqn. (10) we have :

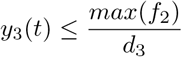

where

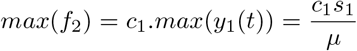

from (14) and therefore

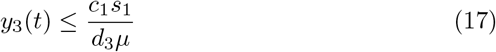
4. Removing the term ‘−*d*_6_*y*_5_(*t*)*y*_4_(*t*)’ from Eqn. (11) since *d*_6_, *y*_5_(*t*) > 0, we can write:

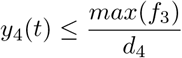

using the fact that *max*(*f*_3_) = *s*_2_ + *p*_3_ we have the boundedness condition on *y*_4_(*t*) as

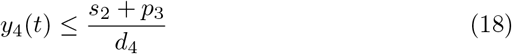
5. Following the same and using (17) we have :

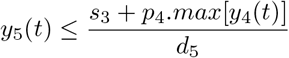

thus

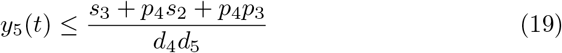 Therefore the boundedness condition for the system from (13),(14),(17),(18) and (19) can be summarised as:

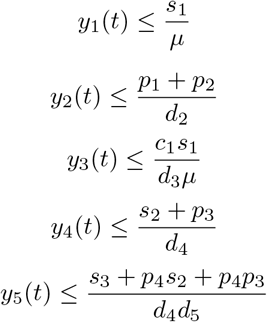

with 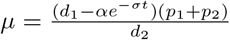 where *d*_1_ > *α* is the required constraint from (15) and (16).

### 3.2 Numerical Results

Numerical simulations were carried out in order to study the model in detail. We first took the sub-model I and on the basis of analysis (see section 3.1.1) we simulated the 3-D model choosing the parameters as required by previously obtained conditions. Table 1 represents the basal set of values of parameters at which the system showed oscillation in both p53 and Mdm2 levels. These oscillations depict the biological scenario of DNA repair process when a stress acts on the system. Next we took this parameter set and did a sensitivity analysis, where we saw the fold range of each parameter around its value in the basal set, keeping other parameters fixed, for the p53 response to be oscillatory. Figure 4 represents the results of the sensitivity analysis, showing that the parameters *α*, *d*_1_ and *s*_1_ were the most sensitive parameters. Now in order to follow the hypothesis as mentioned in section 3.1.1, i.e to have the final state equivalent to the initial state of the system, it is seen that the equality condition *s*_1_ = *d*_1_*y*_10_*y*_20_ should hold true at all times. Keeping this in mind we performed a two parameter study (where the 3rd parameter *s*_1_ is dependent and also varies, but is not shown here explicitly). Figure 5 represents this simulation result, where red region shows the 2-D parameter space of *d*_1_ and *α* at which the oscillations occur and the black region is the space where a high p53 pulse is seen which denotes the phenomena where this 3-D model is unable to repair the DNA damage and marks an initiation of senescence or apoptosis pathways. Figure 6 shows the p53 dynamics at a point in the red oscillatory region and the dynamics when a choice is made from the black space.

**Figure 5:**
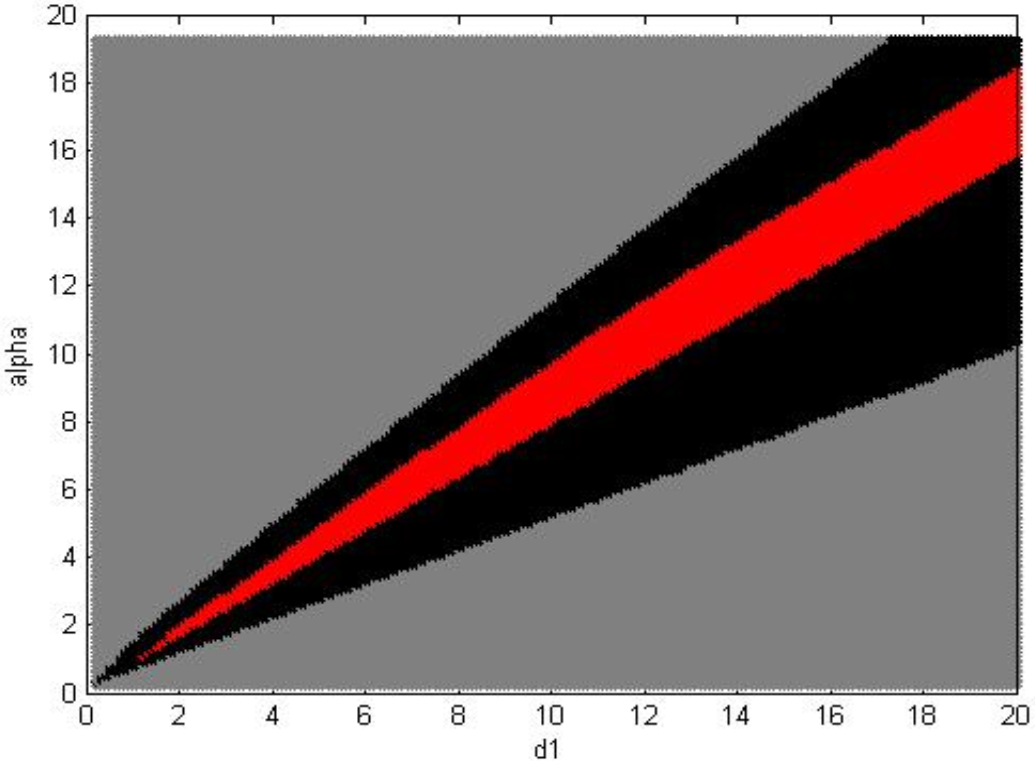
The diagram represents the parameter space of *d*_1_ and *α* where oscillations take place (red) and the area where high p53 pulses appear as model output owing to the phenomenon that there is no DNA repair when this happens (black). The grey color represents non feasible regions in terms of the p53 levels. The upper grey is a region where unbounded p53 levels are seen (violation of condition 16), while the lower grey signifies negligible levels of p53 since input stress is not significant enough (*α* ≪ *d*_1_).

**Figure 6:**
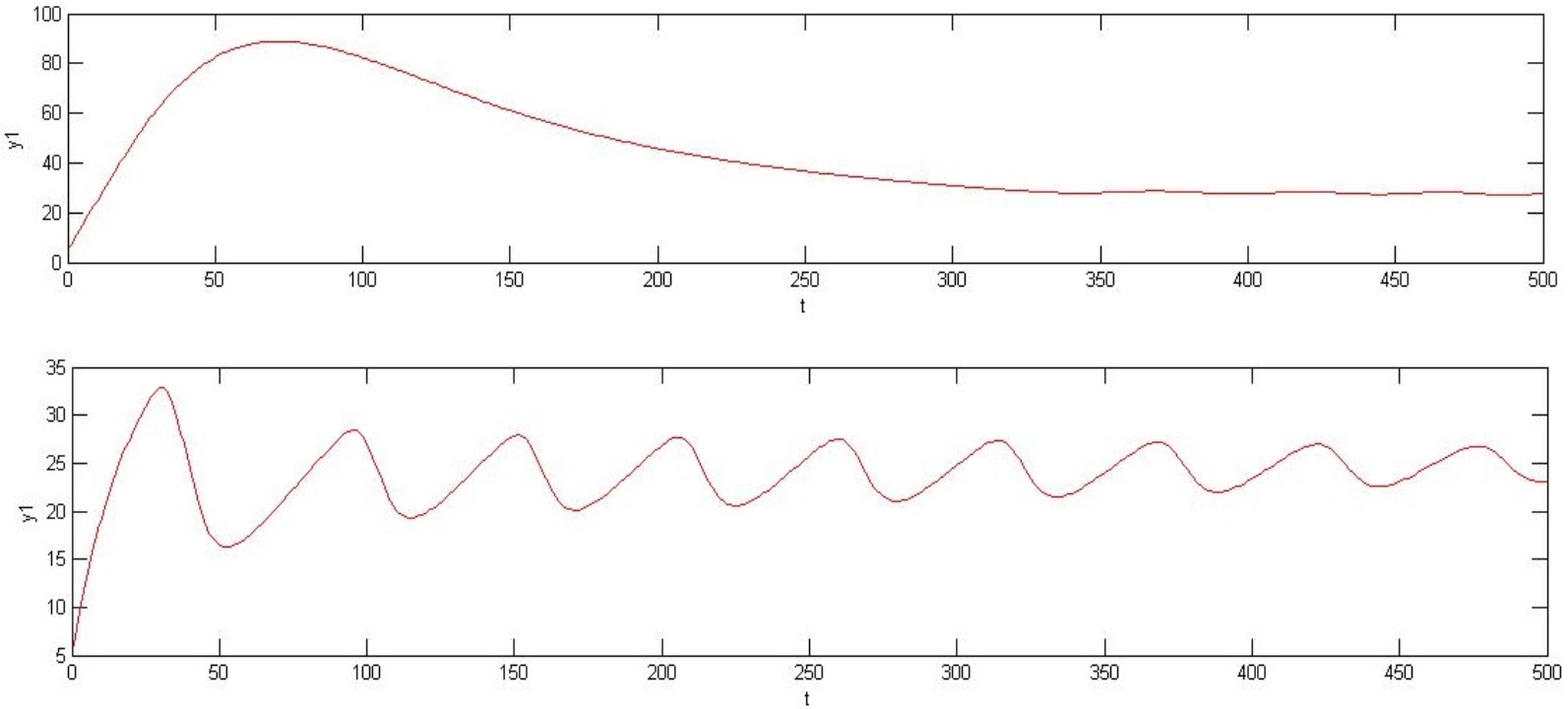
The figure shows the p53 dynamics in time when the *d*_1_, selection is made from the black space (upper graph with *d*_1_ = 8 and *α* = 7.9); and from the red space (upper graph with *d*_1_ = 8 and *α* = 7). All other parameters were fixed at the basal set given in Table 1.

By selecting a set of parameter values from the oscillatory region we then looked at the effect of other parameters, one by one on the system output. The Table 2 shows this result.

**Table 2:** The table shows the range of values of the parameters at *d*_1_ = 8 and *α* = 7. Each parameter except *d*_1_ and *α* is varied one by one, around the basal set as given in Table 1. The effect of parameter variation resulted into either oscillatory response in p53 (Repair) or an over expression of p53 seen as a high pulse (No repair).

| Parameter | Range | Effect |
| --- | --- | --- |
| $c_1$ | $0 - 0.0585$ | No Repair |
| $c_1$ | $0.0585 - 0.21$ | Repair |
| $c_2$ | $0 - 0.25$ | Repair |
| $c_2$ | $> 0.25$ | No Repair |
| $d_3$ | $0.05 - 0.14$ | Repair |
| $d_3$ | $> 0.14$ | No Repair |
| $km_2$ | $11.83 - 39.03$ | Repair |
| $km_2$ | $> 39.03$ | No Repair |
| $n_1$ | $0 - 6$ | No Repair |
| $n_1$ | $> 6$ | Repair |

Next we considered the sub-model II; on the basis of analysis in sec 3.1.2 and using the results above we were able to choose a parameter set (table 3) such that the simulation of the complete model provides us with the biologically relevant scenario viz. (i) When the DNA damage could be repaired i.e when there are oscillations in p53 levels, the senescence pathway is not activated and the IL6 level remain low where as (ii) When there is no possibility of DNA repair i.e no p53 oscillations occur and rather a huge p53 pulse is generated, the cellular senescence pathway is activated and results into high levels of IL6. We show this result in figure 7.

**Table 3:**
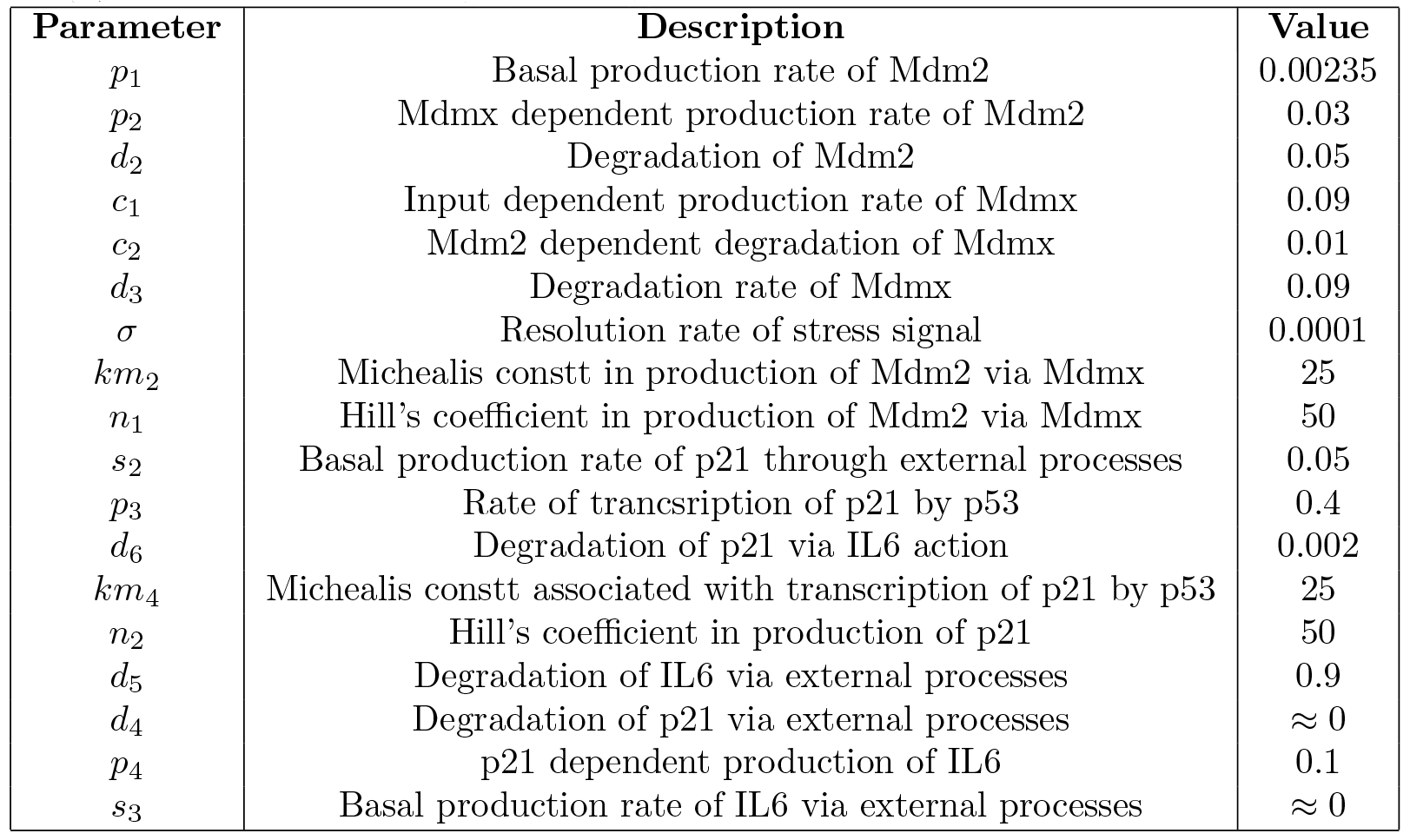
Basal set of parameter values for the complete model to mimic the typical IL6 response in stressed and non stressed conditions. Here initial conditions are *y*_10_ = 5.3191, *y*_20_ = 0.047, *y*_30_ = 2, *y*_40_ = 2 and *y*_50_ = 1.46.

**Figure 7:**
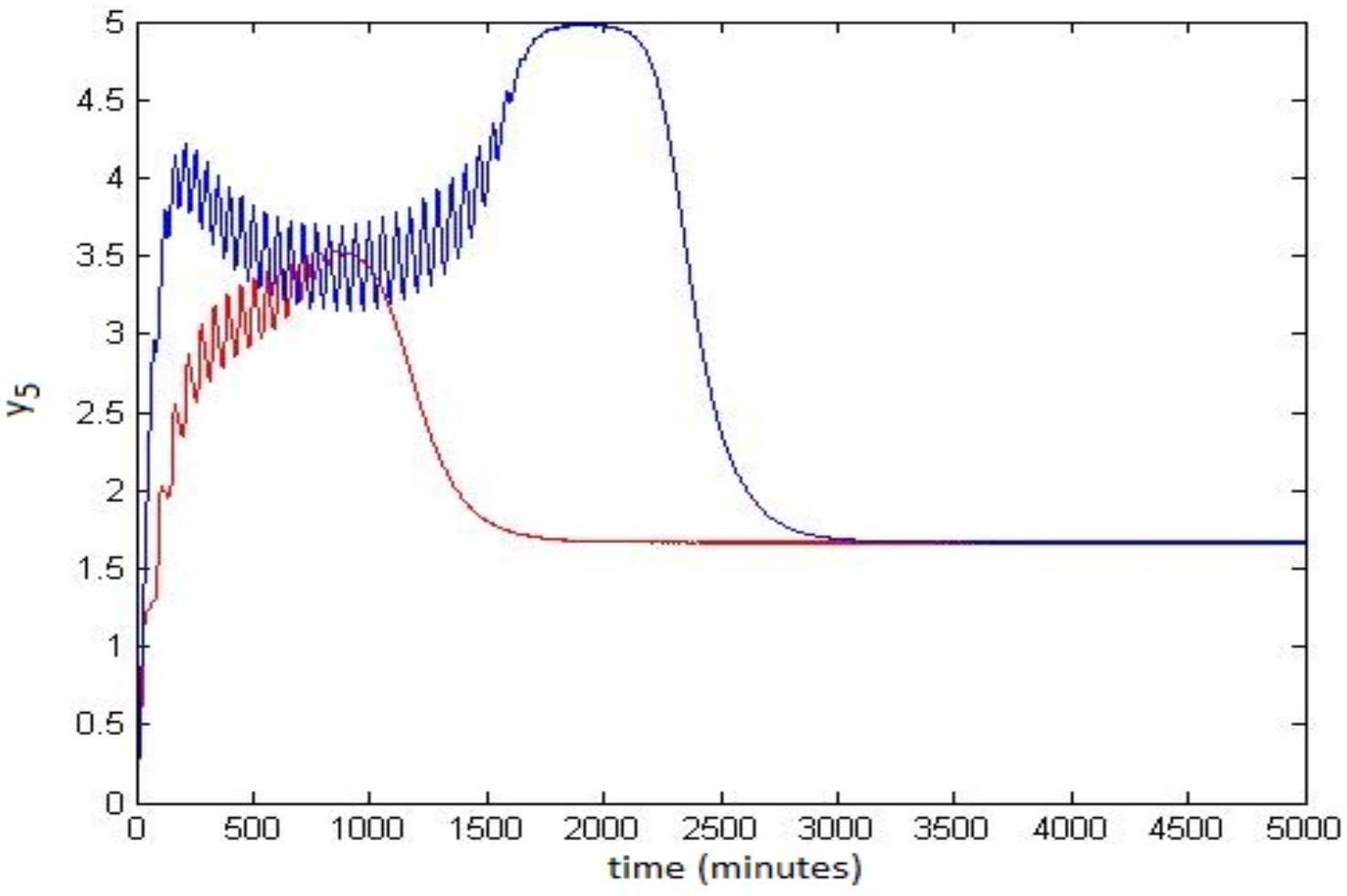
The figure shows the IL6 dynamics under non stressed conditions (red) when *d*_1_ = 8 and *α* = 7 and under stressed conditions (blue) when *d*_1_ = 8 and *α* = 7.9. All the other parameter values are as given in Table 3. Note that the time for which the IL6 levels stay high in the stressed conditions (> 4) is around 15 hours.

The time for which IL6 levels remain high is a crucial quantity for our model in order to form a link between insulin resistance and IL6 levels. A greater time for which IL6 levels remain high would promote insulin resistance to a larger degree. Thus, now we look at the ‘Time for which IL6 levels is higher than a threshold’ v/s the variation in other model parameters (see Figure 8). This threshold is decided on the basis of earlier studies accounting for the fold change of IL6 levels seen in subjects with diabetes type 2 like disease in comparison to normal subjects [38].

**Figure 8:**
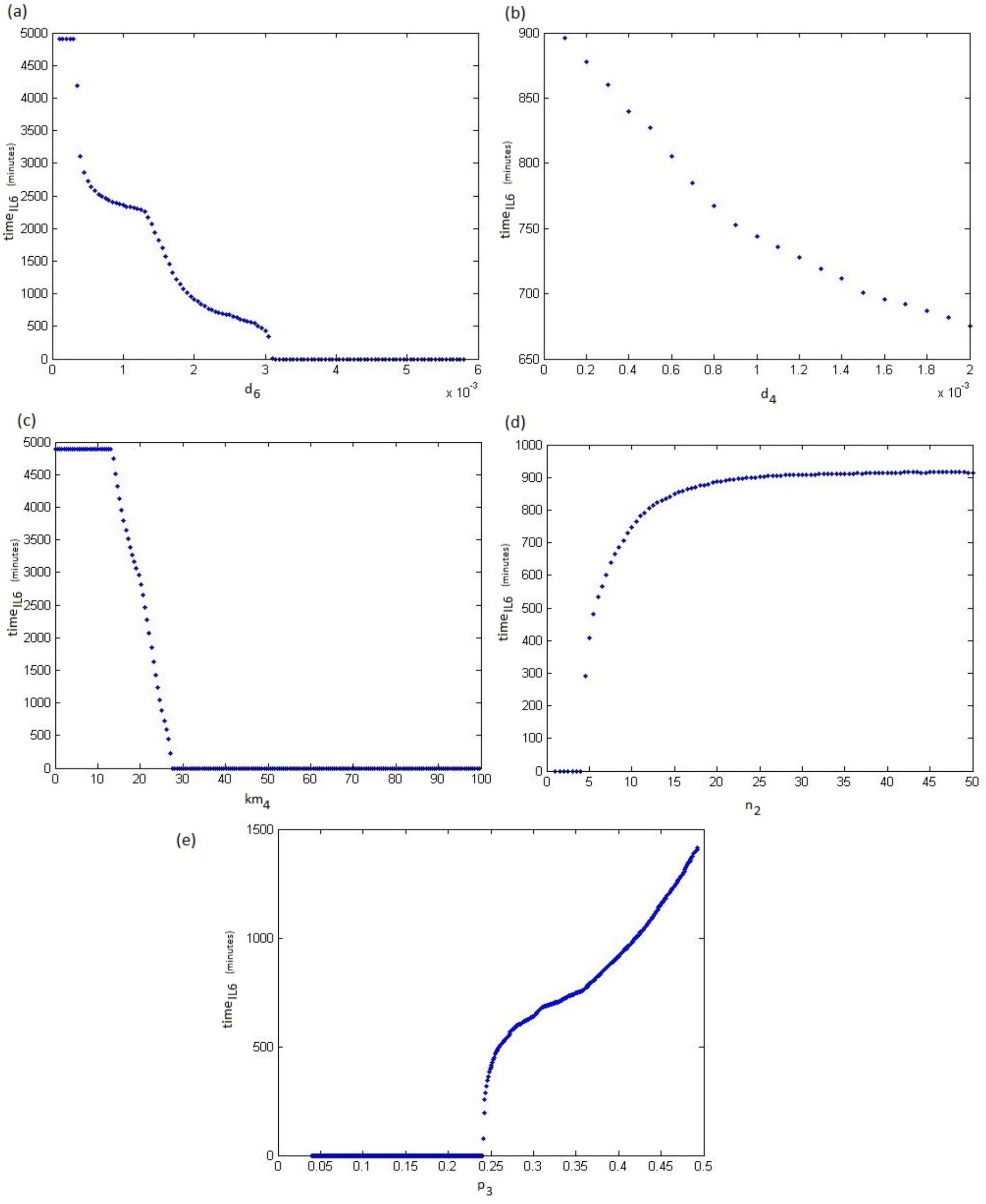
The figure shows variation of IL6 levels with different parameters. The IL6 levels here are quantified in terms of the quantity *time*_*IL*6_ which denotes the time period for which IL6 levels remain high above a basal threshold.

## 4 Discussion

In this study, we have explored the role of different molecular mediators and their interplay in the development of cellular senescence and exacerbation of insulin resistance, in the setting of metabolic dysfunction. Ageing acts as both a risk factor for and an epiphenomenon of insulin resistance, and fuels a vicious cycle of molecular as well as phenotypic perturbations such as chronic low-grade systemic inflammation [44, 45]. Within the milieu of molecules involved in driving metabolic stress-induced premature ageing, our study focused on analysis and modelling of dynamics of the important player p53 and the downstream pathway it triggers towards development of a pro-inflammatory state, a condition fertile for insulin resistance to develop and progress.

Our model showed the existence of oscillations in p53 and Mdm2 levels and in addition, identified the basal rates of the processes involved in maintaining this oscillatory nature (Figure 3 and Table 1).The dynamics of p53 and Mdm2 during normal cell cycle progression have been validated experimentally, where high temporal resolution analysis of p53-Mdm2 levels showed remarkably regulated oscillations [46]. This represents the process of DNA damage and consequent activation of p53 through proteins such as ATM which phosphorylate p53, which triggers repair of damaged DNA and is followed by degradation of p53 by Mdm2. The transient but repetitive nature of this phenomenon, seen in normally growing cells in which DNA damage in low amounts/frequency occurs during the progression of cell cycle, lends oscillatory nature to the dynamics of p53 and Mdm2 levels [46]. Our model showed a stress-induced sharp elevation in p53 levels, as seen in Figure 3 (results). Cellular encounter with a severe genotoxic stress, which could be due to metabolic perturbations and elevated ROS levels, causes a prolonged surge in the levels of p53 that drives cells into growth arrest or premature senescence [8]. Hence, we have mathematically validated the oscillatory dynamics of p53-Mdm2 loop, which is critical in maintaining genomic integrity.

To add to this mathematical recapitulation of cellular senescence under metabolic stress, assessing the degree of effect that various mediators and processes have on the p53 levels was deemed necessary. This was achieved using a sensitivity analysis, whereby we assessed the degree of effect of different conditions on DNA repair through p53, thus identifying processes and conditions that serve as significant determinants of the fate of a cell. For p53 levels to show oscillatory pattern with time (indicating DNA repair), the critical variables identified were the basal production rate of p53 (*s*_1_), stress-induced enhancement in p53 levels (α), and the degradation and inhibition of p53 by Mdm2 (*d*_1_). This finding reflects the high sensitivity of the p53-Mdm2 loop itself, a cellular strategy tipping the critical balance between DNA repair and growth arrest, hence flexibility to constant rewiring being most desirable. Since p53 acts as a powerful suppressor of cell cycle progression, its basal level in cells is maintained under strict control, with only transient pulses seen in response to routine DNA damage occurring during cell cycle [46]. Strong genotoxic stress (ROS generation due to excessive lipolysis in insulin-resistant adipose tissue, for example) induces abruptly high p53 levels and activation, which then triggers the p21-mediated senescent phenotype marked by inflammatory cytokine secretion [8]. Transcriptional activation of Mdm2 by p53 lends self-limiting nature to this process as Mdm2 inhibits as well as degrades p53, and also limits its nuclear localisation. That the sensitivity of these mediators and their cross-talk holds potential for therapeutic modulation of premature cellular senescence in cardio-metabolic disorders is undeniable; our findings in this regard were substantiated by experimental data where statins have been shown to reduce p53 response to DNA damage in stressed conditions by phosphorylating Mdm2 through activation of Akt, and stabilising its negative effect on p53 levels and activity [48].

Once we worked out the requisite dynamics of p53-Mdm2 crosstalk for DNA repair to occur, our focus shifted to the other players in our model as to how they individually affect the maintenance of genomic integrity. Our assumption here was that any variable/factor that restores the oscillatory nature of p53 levels, after a stress acting on the system has led to a p53 surge, would lead to DNA repair and reduce progression to permanent growth arrest [8, 46]. In this analysis, the important dynamics captured were those of Mdmx, the homologue of Mdm2 that inhibits p53 on its own as well as by enhancing stabilisation of Mdm2 [13, 49]. We observed that when Mdmx is at low levels in the cell, p53 levels remain high enough to cause growth arrest and hence repair of DNA is not possible. Beyond certain threshold level of Mdmx (parameter c1 in Table 2.), DNA repair (oscillations in p53 level) was observed, which could be accounted for by the inhibitory interaction of Mdmx with p53 and also its positive effect on Mdm2 stability. Proteasomal degradation of Mdmx occurs after ubiquitination by Mdm2; phosphorylation of S367 residue in response to DNA damage by certain proteins such as ATM and Chk1 has also been shown to cause degradation of Mdmx [28]. Our results clearly showed that when degradation of Mdmx is beyond a certain threshold (parameter *d*_3_ in Table 2), DNA repair is possible and that, crossing this critical rate of degradation makes the cell progress to premature senescence. This finding is biologically plausible since many studies have shown that Mdmx is phosphorylated and degraded following DNA damage, allowing a rapid build-up of p53 levels in the cell. An interaction with c-Abl downstream of ATM has been shown to phosphorylate Mdmx at Y99, ultimately interfering with its interaction with p53, eventually activating p53 [51]. A desirable strategy to favourably manoeuvre cellular fate away from senescent phenotype would thus be manipulating Mdmx such that p53 levels and activation in response to stress could be controlled. According to a study, insulin-like growth factor-1 (IGF1) can increase levels of Mdmx mRNA and subsequently its protein levels [51]. Also, Akt was found to directly phosphorylate Mdmx at S367, enhancing 14-3-3 binding, which actually stabilized Mdmx and downregulated p53 [52]. This finding echoes the results of another study that show Akt-activating drug pravastatin can rescue biological system from stress-induced p53 activation, and provides a lead for the potential management of DNA damage-induced senescence and subsequent exacerbation of insulin resistance [48].

Our study was meant to mathematically analyse the scenario of premature cellular ageing in metabolic syndrome and its complications, involving the development of senescence-associated secretory phenotype (SASP), which is marked by elevated levels of pro-inflammatory cytokines such as IL-6, TNF-a, IL-1 etc. After analysis of the p53-Mdm2 component, we chose to study the dynamics of IL-6 that forms a crucial link in the chain of progression from DNA damage-induced p53-p21 axis of senescence to development and progression of insulin resistance. Il-6 is a pleiotropic cytokine responsible for inflammatory response that could be both acute, and chronic as in case of insulin resistance. Our model showed stress-induced prolonged elevation of IL-6 level, supported by stress-induced elevation of IL-6 via p53-p21 (WAF) axis being well-documented as experimental evidence. An interesting finding of our analysis was the temporal pattern of more than two-fold elevation in IL-6 levels (threshold decided on the basis of published clinical evidence) [38], where we saw that IL-6 levels stay high for about fifteen hours after input stress acts on the system. Making this observation in-vivo i.e. in animal models of metabolic syndrome or in obese/insulin-resistant individuals would have been tough since the frequency of impulse (stress), which would play a major role on the time-profile of IL-6, is difficult to determine or manipulate in a living system where each regulatory circuit is embedded over a background of many others. Nevertheless, we can clearly see the impact of stress on nature of elevation in IL-6 levels and fathom the biological implications of this effect where presence of persistent stress causes development of chronic inflammatory phenotype both at systemic and tissue level in individuals with metabolic syndrome [53].

An important objective of this study was to figure out possible strategies to manage the inflammatory state associated with senescence, without fiddling with p53-Mdm2 dynamics directly. The reason was the highly critical role of p53-Mdm2 interaction in cell cycle regulation where senescence itself acts like a double-edged sword due to growth arrest leading to undesirable premature ageing and associated complications, but preventing escape of cells from normal cell cycle to cancerous character. Hence, we analysed the effect of different conditions/processes (system parameters) on the temporal dynamics of IL-6 levels in order to identify candidate molecules/interactions that could serve as targets for therapeutic intervention. The critical determinants of IL-6 time-profile, as identified from our analysis, revolved around p21 viz. p53-mediated transcription and, degradation/inhibition through Akt as well as other sources. That p53-induced transcription of p21 shows a steep rise but only after a threshold was seen in our findings (Figure 8(e)). This could be explained by the findings where constitutive/basal levels of p53 during cell cycle progression did not induce p21 transcription even when it is activated in response to routine DNA damage; severe challenge posed to DNA integrity by DNA-damaging treatment such as UV irradiation or oxidative insult induces p53-guided transcriptional activation of p21 [46]. Identifying this cellular threshold in clinical setting is not an easy task; lifestyle and pharmacological approaches that minimise the metabolic stress are proven to be desirable nevertheless. Further findings from our models analysis involving IL-6 added to the evidence supporting the positive role of Akt in senescence, and its contribution to resolving inflammatory phenotype. Akt has been shown to induce cytoplasmic localisation of p21 through phosphorylation, ultimately reducing the activity of p21 in the growth arrest pathway [54], which is a plausible explanation for our results shown in Figure 8(a). Similar shortening of the duration of IL-6 elevation was seen on account of p21 inhibition/degradation by other mechanisms (Figure 8(b)).

Our study pointed toward modulation of the metabolic stress-senescence-insulin resistance axis at multiple levels, the master regulator for which appeared to be Akt. The mechanisms acting in ageing and associated disorders are intricately woven and not fully understood. In such a scenario, proven reciprocal relationship of Akt with the premature ageing p53-p21 axis at so many levels raises expectations about Akt-activating drugs in ameliorating insulin resistance accounted for by senescence-induced inflammatory state. This strategy seems particularly lucrative considering the established status of Akt as a regulator of metabolism whose activation is linked with reversal of insulin resistance through pathways other than p53-p21-IL6 axis.

Our findings raise some interesting observations and questions regarding the contribution of premature ageing to insulin resistance and possible strategies for its management. Amelioration of elevated IL-6 levels through inhibition of p21, a strategy meant to rescue cells from associated insulin resistance with minimal interference in the tumour-suppressor role of p53, may indirectly itself take care of cellular fate as chronic inflammation due to IL-6 has also been associated with carcinogenesis [56, 57]. Also, exploring the temporal dynamics of IL-6 elevation due to metabolic stress, and its association with insulin resistance would be an exciting arena if validated by experimental biologists, with publications relating time-profile of action to nature of effect of cytokines already coming up [58, 59].

## 5 Conclusion

We formulated a mathematical model for metabolic stress-induced pro-inflammatory state mediated by activation of p53 and downstream mediators. Since the aim was to find a strategy to rejuvenate ageing cells, we identified the basal conditions required for efficient DNA repair, and also graded the effect of each on the system using sensitivity analysis. Our model validated the remarkable flexibility of p53-Mdm2 loop for efficient adaptation to varying cellular conditions and environmental stimuli, and the role of Mdmx in modulating this interplay. We modeled the hitherto unreported temporal dynamics of IL-6 by elucidating, to a great degree, the effect of members of the p53-p21 senescence axis on the time profile of its elevation. Our results provide a lead for possible therapeutic strategies to curb chronic inflammation and eventually insulin resistance through activation of Akt, without disturbing the p53-Mdm2 loop directly, thus preventing possible interference in cancer-suppression mechanisms. Overall, this study adds a new dimension to the field finding strategies aimed at management of ageing-associated insulin metabolic complications.

## References

[1] Global status report on noncommunicable diseases 2014. Geneva, WHO.

[2] World Health Organization, (2014). Global Health Estimates: Deaths by Cause, Age, Sex and Country, 2000-2012. Geneva, WHO.

[3] Mathers, C.D., Loncar, D. (2006). Projections of global mortality and burden of disease from 2002 to 2030. PLoS Med. 3(11):e442.

[4] Monk, M., (2003). Oscillatory expression of Hes1, p53, and NF-kB driven by transcriptional time delays. Current Biology. 13:14091413.

[5] Qi, J.P., et al., (2007). A mathematical model of P53 gene regulatory networks under radiotherapy. BioSystems. 90:698706

[6] Anderson, A.R.A., (2005). A hybrid mathematical model of solid tumour invasion: the importance of cell adhesion. Mathematical Medicine and Biology. 22:163186

[7] Simone, S., et al., (2008). “Mechanism of oxidative DNA damage in diabetes tuberin inactivation and downregulation of DNA repair enzyme 8-oxo-7, 8-dihydro-2’-deoxyguanosine-DNA glycosylase.” Diabetes. 57.10:2626–2636.

[8] Monickaraj, F., et al., (2013). Accelerated fat cell aging links oxidative stress and insulin resistance in adipocytes. J. Biosci. 38(1):113–22.

[9] Minamino, T., et al., (2009). A crucial role for adipose tissue p53 in the regulation of insulin resistance. Nature Medicine. 15.9:1082–1087

[10] Homayounfar, R., et al., (2015). Relationship of p53 accumulation in peripheral tissues of high-fat diet-induced obese rats with decrease in metabolic and oncogenic signaling of insulin. General and comparative endocrinology. 214:134–139.

[11] Geva-Zatorsky, et al., (2006). Oscillations and variability in the p53 system. Mol. Syst. Biol. 2:.0033.

[12] Bar-Or, R.L., et al., (2000). Generation of oscillations by the p53-Mdm2 feedback loop: a theoretical and experimental study. Proc. Natl. Acad. Sci. USA 97, 1125011255.

[13] Stad, R., et al., Hdmx stabilizes Mdm2 and p53. J. Biol. Chem. 2000;275:28039–44.

[14] Hasty, P., Christy, B. A., (2013). p53 as an intervention target for cancer and aging. Pathobiology of aging and age related diseases. 3.

[15] de Oca Luna, R. M., et al., (1995). Rescue of early embryonic lethality in mdm2-deficient mice by deletion of p53. Nature. 378(6553):203–206.

[16] Oliner, J. D., et al., (1992). Amplification of a gene encoding a p53-associated protein in human sarcomas. Nature. 358.

[17] Freedman, D. A., et al., (1999). Functions of the Mdm2 oncoprotein. Cellular and Molecular Life Sciences. 55(1):96–107.

[18] Juven-Gershon, T., Oren, M., (1999). Mdm2: the ups and downs. Molecular medicine. 5(2):71.

[19] Kubbutat, M. H., et al., (1997). Regulation of p53 stability by Mdm2. Nature. 387(6630):299–303.

[20] Haupt, Y., et al., (1997). Mdm2 promotes the rapid degradation of p53. Nature. 387(6630):296–299.

[21] Bttger, A., et al., (1997). Design of a synthetic Mdm2-binding mini protein that activates the p53 response in vivo. Current Biology. 7(11):860–869.

[22] Honda, R., Yasuda, H. (1999). Association of p19ARF with Mdm2 inhibits ubiquitin ligase activity of Mdm2 for tumor suppressor p53. The EMBO journal. 18(1):22–27

[23] Damalas, A., et al., (2001). Deregulated-catenin induces a p53-and ARF-dependent growth arrest and cooperates with Ras in transformation. The EMBO journal. 20(17):4912–4922.

[24] Barak, Y., et al., (1994). Regulation of mdm2 expression by p53: alternative promoters produce transcripts with nonidentical translation potential. Genes and development. 8(15):1739–1749.

[25] Saucedo, L. J., et al., (1998). Regulation of transcriptional activation of mdm2 gene by p53 in response to UV radiation. Cell growth and differentiation. 9(2):119–130.

[26] Kawai, H., et al., (2007). RING DomainMediated Interaction Is a Requirement for Mdm2’s E3 Ligase Activity. Cancer Research. 67(13):6026–6030.

[27] Linares, L. K., et al., (2003). HdmX stimulates Hdm2-mediated ubiquitination and degradation of p53. Proceedings of the National Academy of Sciences. 100(21):12009–12014.

[28] Pereg, Y., et al., (2006). Differential roles of ATM-and Chk2-mediated phosphorylations of Hdmx in response to DNA damage. Molecular and cellular biology. 26(18):6819–6831.

[29] Shvarts, A., et al., (1997). Isolation and identification of the human homolog of a new p53-binding protein, Mdmx. Genomics. 43(1):34–42.

[30] Passos, J. F., et al., (2010). Feedback between p21 and reactive oxygen production is necessary for cell senescence. Molecular systems biology. 6(1):347.

[31] Hotamisligil, G. S., et al., (1995). Increased adipose tissue expression of tumor necrosis factor-alpha in human obesity and insulin resistance. Journal of Clinical Investigation. 95(5):2409.

[32] Bastard, J. P., et al., (2002). Adipose tissue IL-6 content correlates with resistance to insulin activation of glucose uptake both in vivo and in vitro. The Journal of Clinical Endocrinology and Metabolism. 87(5):2084–2089.

[33] Kojima, H., et al., (2013). IL-6-STAT3 signaling and premature senescence. JAK-STAT. 2(4):e25763.

[34] Li, Y., et al., (2002). AKT/PKB phosphorylation of p21Cip/WAF1 enhances protein stability of p21Cip/WAF1 and promotes cell survival. Journal of Biological Chemistry. 277(13):11352–11361.

[35] Jeong, J. H., et al., (2010). p53-independent induction of G1 arrest and p21WAF1/CIP1 expression by ascofuranone, an isoprenoid antibiotic, through downregulation of c-Myc. Molecular cancer therapeutics. 9(7):2102–2113.

[36] Sankala, H. M., (2007). Involvement of sphingosine kinase 2 in p53-independent induction of p21 by the chemotherapeutic drug doxorubicin. Cancer research. 67(21):10466–10474.

[37] Kishimoto, T., (2010). IL-6: from its discovery to clinical applications. International immunology. 22(5):347–352.

[38] Kern, P. A., et al., (2001). Adipose tissue tumor necrosis factor and interleukin-6 expression in human obesity and insulin resistance. American Journal of Physiology-Endocrinology And Metabolism. 280(5):E745–E751.

[39] Ciliberto, A., et al., (2005). Steady states and oscillations in the p53/Mdm2 network. Cell cycle. 4(3):488–493.

[40] Novk, B., Tyson, J. J., (2008). Design principles of biochemical oscillators. Nature reviews Molecular cell biology. 9(12):981–991.

[41] Tyson, J. J., et al., (2003). Sniffers, buzzers, toggles and blinkers: dynamics of regulatory and signaling pathways in the cell. Current opinion in cell biology. 15(2):221–231.

[42] Gouz, J. L., (1998). Positive and negative circuits in dynamical systems. Journal of Biological Systems. 6(01):11–15.

[43] Brooks, C. L., Gu, W., (2006). p53 ubiquitination: Mdm2 and beyond. Molecular cell. 21(3):307–315.

[44] Dubowitz, N., et al., (2014). Aging is associated with increased HbA1c levels, independently of glucose levels and insulin resistance, and also with decreased HbA1c diagnostic specificity. Diabetic Medicine. 31(8):927–935.

[45] Law, I. K., et al., (2010). Lipocalin-2 deficiency attenuates insulin resistance associated with aging and obesity. Diabetes. 59(4):872–882.

[46] Loewer, A., et al., (2010). Basal dynamics of p53 reveal transcriptionally attenuated pulses in cycling cells. Cell. 142(1):89–100.

[47] Rodier, F., Campisi, J., (2011). Four faces of cellular senescence. The Journal of cell biology. 192(4):547–556.

[48] Paajarvi, G., et al., (2005). HMG-CoA reductase inhibitors, statins, induce phosphorylation of Mdm2 and attenuate the p53 response to DNA damage. The FASEB journal. 19(3):476–478.

[49] Sharp, D. A., et al., (1999). Stabilization of the Mdm2 oncoprotein by interaction with the structurally related Mdmx protein. Journal of Biological Chemistry. 274(53):38189–38196.

[50] Zuckerman, V., et al, (2009). c-Abl phosphorylates Hdmx and regulates its interaction with p53. Journal of Biological Chemistry. 284(6):4031–4039.

[51] Gilkes, D. M., et al., (2008). Regulation of Mdmx expression by mitogenic signaling. Molecular and cellular biology. 28(6):1999–2010.

[52] Lopez-Pajares, et al., (2008). Phosphorylation of Mdmx mediated by Akt leads to stabilization and induces 14-3-3 binding. Journal of Biological Chemistry. 283(20):13707–13713.

[53] Rotter, V., et al., (2003). Interleukin-6 (IL-6) induces insulin resistance in 3T3-L1 adipocytes and is, like IL-8 and tumor necrosis factor-a, overexpressed in human fat cells from insulin-resistant subjects. Journal of Biological Chemistry. 278(46):45777–45784.

[54] Zhou, B. P., et al., (2001). Cytoplasmic localization of p21Cip1/WAF1 by Akt-induced phosphorylation in HER-2/neu-overexpressing cells. Nature Cell Biology. 3(3):245–252.

[55] Ota, H., et al., (2010). Induction of endothelial nitric oxide synthase, SIRT1, and catalase by statins inhibits endothelial senescence through the Akt pathway. Arteriosclerosis, thrombosis, and vascular biology. 30(11):2205–2211.

[56] Nguyen, D. P., et al., (2014). Inflammation and prostate cancer: the role of interleukin 6 (IL-6). BJU international. 113(6):986–992.

[57] Taniguchi, K., Karin, M., (2014). IL-6 and related cytokines as the critical lynchpins between inflammation and cancer. Seminars in immunology 26(1):54–74.

[58] Braun, D. A., et al., (2013). Cytokine response is determined by duration of receptor and signal transducers and activators of transcription 3 (STAT3) activation. Journal of Biological Chemistry. 288(5):2986–2993.

[59] Raschzok, N., et al., (2011). Temporal expression profiles indicate a primary function for microRNA during the peak of DNA replication after rat partial hepatectomy. American Journal of Physiology-Regulatory, Integrative and Comparative Physiology. 300(6):R1363–R1372.

[60] Vousden, K. H., Prives, C., (2009). Blinded by the light: the growing complexity of p53. Cell. 137(3):413–431.

[61] Appella, E., Anderson, C.W. (2001). Post-translational modifications and activation of p53 by genotoxic stresses. Eur. J. Biochem. 268:2764–2772

[62] Augustyn, K.E., et al., (2007). A role for DNA-mediated charge transport in regulating p53: Oxidation of the DNA-bound protein from a distance. Proc. Natl. Acad. Sci. 104:18907–18912

[63] Bensaad, K., Vousden, K.H., (2007). p53: new roles in metabolism. Trends Cell Biol. 17:286–291.

